# Plant-derived insulator-like sequences for control of transgene expression

**DOI:** 10.1101/2021.11.04.467280

**Authors:** Jubilee Y. Park, Lynsey Kovar, Peter R. LaFayette, Jason G. Wallace, Wayne A. Parrott

## Abstract

Stable and consistent transgene expression is necessary to advance plant biotechnology. Stable expression can be achieved by incorporating enhancer-blocking insulators, which are *cis*-regulatory elements that reduce enhancer interference in gene expression, into transgene constructs. Sufficient insulators for plant use are not available, and their discovery has remained elusive. In this work, we computationally mined the compact genome of *Utricularia gibba* for insulator sequences and identified short (<1 kb) sequences with potential insulator activity. Based on *in vivo* tests, three of these effectively mitigate the ectopic transgene expression caused by the Cauliflower Mosaic Virus 35S promoter and do so better than previously reported plant insulators. However, all sequences with apparent insulator activity also decrease the effectiveness of the CaMV 35S promoter, and thus may be more accurately classified as silencers. However, since the insulator effect is proportionately much higher than the silencing effect, these sequences are still useful for plant transformation.

## Main

Insulators are cis-regulatory elements that reduce interference in gene expression. Such cross-interference with transgenes is problematic for crop development as the demand for crops with multiple transgenes continues to grow. Of the 191.7 million hectares dedicated to growing genetically engineered crops in 2021, 42% were planted with crops with stacked traits, meaning multiple traits conferred by transgenes [1]. These mostly were obtained by crossing individual transgenics together. Going forward, multiple transgenes would be ideally placed in one locus to facilitate downstream breeding, but such molecular stacks are difficult to achieve while maintaining proper control of gene expression.

Like other organisms, plants are hypothesized to have insulators based on their evolutionarily conserved genome function and organization, and on gene expression patterns that hint of insulator function [2]. Nevertheless, plant insulators lack sequence conservation to known insulators and have been notoriously difficult to identify and isolate. Furthermore, insulators can be confused with silencers. Insulators must be placed between the enhancer and the promoter. In contrast, silencers are another type of element that retains function independently of its orientation or location [3]. Regardless, the lack of compact and dependable insulators is becoming recognized as a limitation for plant biotechnology applications when enhancers are present.

Enhancers are cis-regulatory transcription elements that are bound by transcription factors to increase transcription of a gene and are position and orientation-independent. The presence of enhancers can trigger unwanted enhancer-promoter interactions, thereby disturbing the tissue-specificity and strength of promoters in transgenic plants [4]. Enhancers work to bridge distances by forming chromatin loops to bring the enhancer and promoter gene into proximity [5]. The commonly used plant promoter Cauliflower Mosaic Virus 35S promoter (CaMV 35S) is a prime example of this phenomenon (Figure 1); it has been shown to activate nearby tissue-specific promoters and cause ectopic expression with nearby transgenes, even in genes up to 4.3-kb way from the T-DNA integration site [6]. Insulators have been well-characterized in animals and are classified into two groups: enhancer-blocking insulators, which interfere with enhancer-promoter interactions when placed between them [7], and scaffold/matrix attachment regions (S/MARs), which can protect a flanked transgene by preventing the spread of a heterochromatin state and the gene silencing it causes [8]. Numerous S/MARs have been isolated from plant species [9], but flanking a transgene with these S/MAR sequences does not consistently reduce transgene expression variability [4, 10], and in some cases, it even increases interference [9].

**Fig 1.**
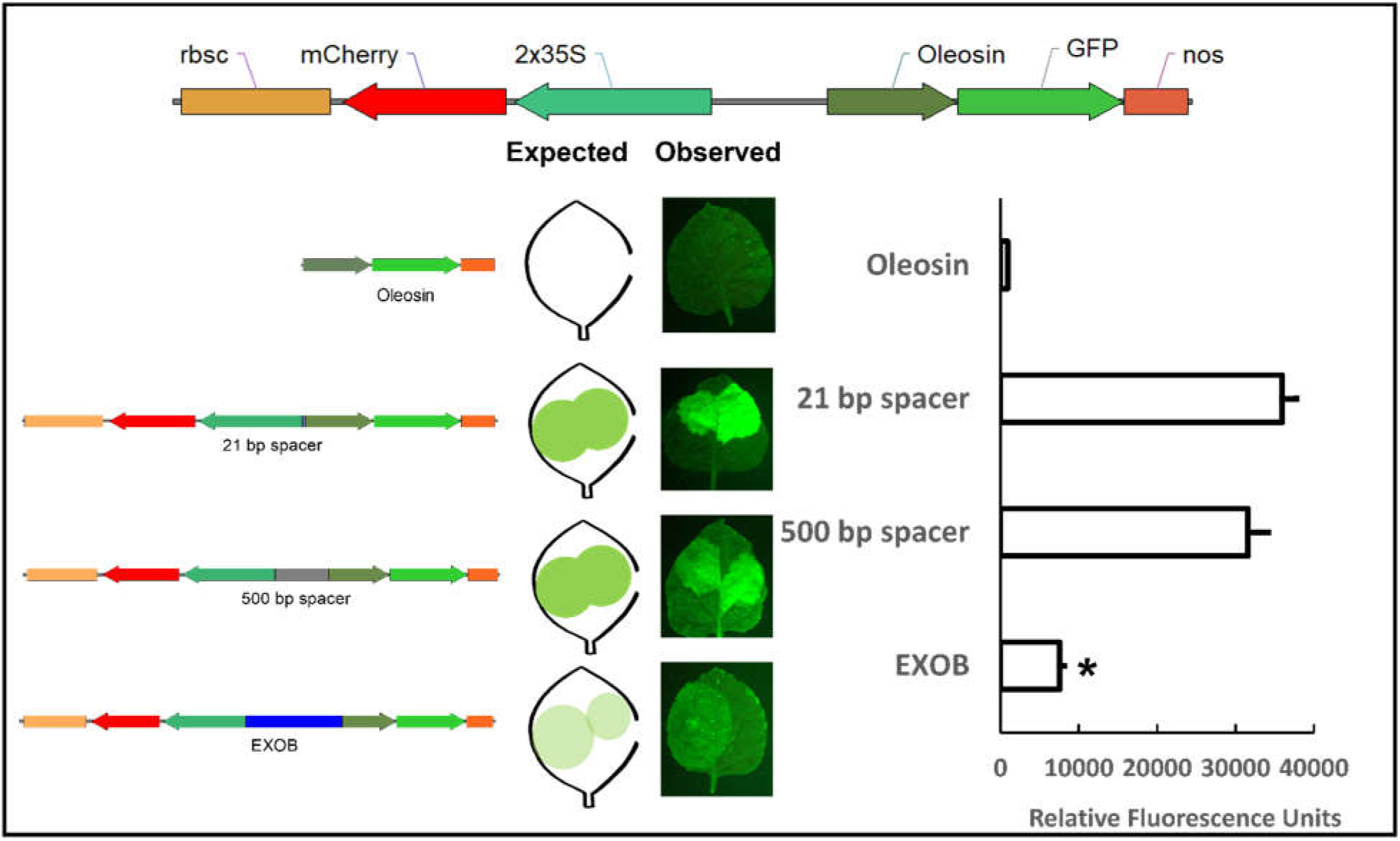
Test construct design, GFP fluorescence in infiltrated *Nicotiana benthamiana* leaves, and representative leaf images. rbsc: rubisco small subunit terminator from *Pisum sativum*, 2×35S: duplicated 35S promoter, Oleosin: *Glycine max* oleosin promoter, nos: Nopaline synthase terminator. Construct names denote the sequence between the 2×35S and Oleosin promoter. Lack of GFP expression is indicative of insulator activity. In this figure, placing the 35S:mCherry cassette in front of oleosin:GFP leads to ectopic expression of GFP. *Indicates t-test significant difference from the control construct (Oleosin promoter with 21 bp between cassettes) p < 0.001. Error bars represent standard error of the mean.

The 2-kb TBS (Transcription Booster Sequence) from *Petunia hybrida* was among the first plant insulator sequences reported based on its effectiveness in both *Arabidopsis* and tobacco [11, 12]. The same group serendipitously discovered EXOB, a 1-kb sequence from phage *Lambda* [11] and found that a 2.2-kb *gypsy*-like sequence from *Arabidopsis* with effective insulator activity [13]. Similarly, Gudynaite-Savitch et al, 2009 [14] also reported that an approximately 900-bp sequence from *Drosophila* (Fab-7 PRE) sequence had insulator function in *Arabidopsis*. However, to be useful in plant transformation, insulators must be as short as possible, as the large size of these insulators limits their usefulness in multigene cassettes by making the cassettes too unwieldy to be readily transformable.

Shorter insulators have been reported, including a 154-bp LTR sequence from the HIV-1 virus [10, 11], and N129, a 16-bp fragment from *Arabidopsi*s [12]. There is a 185-bp element within the HSP70 promoter that is critical for promoter activity but also has some insulator activity [15]. NI29, the 16-bp insulator from *Arabidopsis*, was an ideal and promising insulator because of its small size; however, it does not consistently block the influence of the CaMV 35S promoter, and in some cases, it actually increases ectopic expression [12, 14].

Two other short non-plant-derived insulators, BEAD1c (538 bp) from *Homo sapiens* and UASrpg (234 bp) from the fungus, *Ashbya gossypii*, have demonstrated insulator function in *Arabidopsis* [14]. These partially retain their insulator function when inverted [16], although their insulator ability depends on the genes involved. The use of these insulators in plants has not been studied beyond the initial report.

Sequencing and bioinformatics approaches that have been used to mine plant genomes for novel *cis-*regulatory elements [17, 18] can be extended to insulators. Towards that end, *Utricularia gibba*, or humped bladderwort (Figure 2), is a promising source of insulators. Bladderwort is an aquatic species of carnivorous weed found throughout North America, and it has one of the smallest and most compact genomes among angiosperms: flow cytometry places the genome at ∼88 Mb [19] while a recent genome assembly is ∼102 Mb in length and contains 29,666 genes [20]. The discrepancy could be due to intra-species variation or uncollapsed regions of the genome assembly.

**Fig 2.**
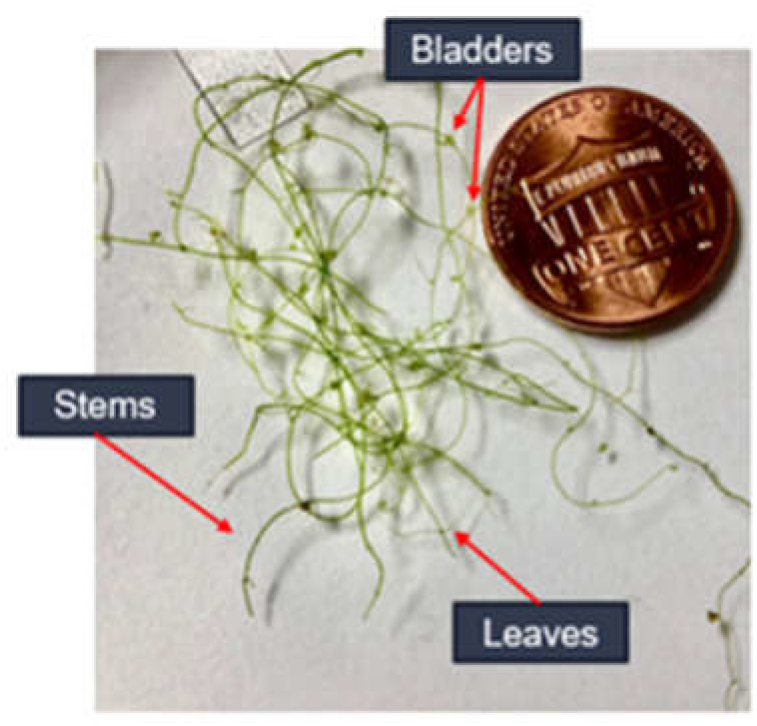
***Utricularia gibba*** stems, bladder, leaves, and growth architecture.

For comparison, the genome of the model species *Arabidopsis* is 135 Mb long and has 33,252 genes [20]. The bladderwort genome contains a smaller-than-average amount of intergenic sequence relative to other sequenced angiosperm genomes. Despite this small amount of intergenic sequence, regulation of genes is tightly controlled, with many genes showing greater than 100-fold differences in expression levels despite being separated by fewer than 1 kb [20].

Bladderwort has comparatively small intergenic sequences, which comprise only 50% of its genome versus 85% in tomato and 95% in maize. From the published PacBio genome assembly, it is evident that 62% of gene pairs are less than 1 kb apart, and of these gene pairs, many show a > 50-fold difference in expression despite being adjacent [21]. All of these traits make bladderwort a promising candidate for mining novel and compact cis-regulatory elements.

The bladderwort genome was computationally mined for putative insulator elements. Bladderwort was re-sequenced, building upon the existing annotation [20]. Three-prime RNA-seq data were used to mine the intergenic regions associated with the presence of insulator elements, producing a ranked list for the top 10 candidate sequences (Table S1). This list was based on several criteria: (1) the magnitude of expression differences between neighboring genes, (2) evidence of independent regulation, and (3) size of the intergenic region, with small regions preferred over large ones.

## Results

### Genome assembly and annotation

*U. gibba* has two public genome sequences, an initial Illumina assembly [21] and a later improved assembly based on PacBio reads [20] from the same accession collected from Michoacán, Mexico. After several attempts to get a sample of this material failed, we obtained live *U. gibba* from a public resource and used Illumina sequencing to create a draft assembly of this specific accession, which we call “PP01”. Although our draft assembly is more fragmented than the published sequence, the final gene count and BUSCO score—a measure of identified conserved genes [22]--are essentially the same (Table S2). This indicates that our assembly covers most of the known gene space.

### Identifying potential insulators

To identify potential insulators, we performed 3’ RNA-seq on four different tissues: rhizoid (a root-like tissue), stem, leaf, and bladder. The resulting gene expression data were combined with the genome annotation to look for putative insulators between gene pairs. Our criteria for flagging intergenic regions as potential insulators depended on the gene orientation of the genes in the pair and expression correlation (or lack thereof) across tissues. We required both genes to have at least some expression to avoid false calls due to misannotations, and at least one gene to be in the top 50% of expressed genes overall to enrich for stronger insulators and avoid calling ones based only on stochastic noise. Putative insulators were flagged based on uncorrelated gene expression between the pair. In total we identified 43 putative insulators passing our thresholds, 16 of which showed conservation across asterids based on whole-genome alignment. We prioritized short elements (<1000 bp) over longer ones as our goal was to identify insulators useful for biotechnology, not to perform an exhaustive annotation of these elements in the genome.

### The test system

To investigate the effectiveness of the 10 identified putative bladderwort insulators, we developed a series of test vectors (Table S3). Each vector contains two reporter genes cassettes: one with GFP driven by the *Glycine max* oleosin promoter, a tissue-specific promoter that is not expressed in *Nicotiana benthamiana* leaves; the second cassette is in the opposite orientation and consists of the duplicated CaMV 35S promoter (2×35S) that drives the expression of the mCherry reporter gene. In control vectors, a minimal amount of spacer DNA (21 bp) separates the two cassettes. The absence of insulator activity results in the enhancer in the 2×35S promoter causing ectopic expression of GFP in tobacco leaves [14] (Figure 1). In other vectors, the cassettes are separated by an approximately 500-bp spacer or by the sequence being tested for insulator activity. Effective insulators will stop enhancer interference from the 2×35S promoter and block GFP expression in leaves. The *U. gibba* insulators were also compared to the HIV-LTR, TBS, and EXOB sequences [11, 23], all of which have been reported to demonstrate insulator activity in plants. Two animal insulators that have been reported to work in plants (BEAD1c and UASrpg) [14, 16] were compared in our test system as well.

### Spacer DNA does not consistently block enhancer interference from the 2×35S promoter

Enhancers are known to work at distances as far away as several Mb [24]. Yet, in some cases, ∼3 kb between the 35S promoter and tissue-specific promoters can dampen ectopic interference [14, 25] in combination with some promoters, perhaps suggesting the presence of a silencer in the sequence. To test this, we compared the difference in ectopic GFP expression between the 21-bp spacer DNA construct and the 500-bp spacer DNA construct, with 500 representing a practical upper size limit. As evident from Figure 1, adding 500 bp between the cassettes did not lead to significant differences in GFP expression (*p*= 0.094).

### Bladderwort insulators

Three standout bladderwort insulators were identified that consistently resulted in little to no GFP expression across three separate trials; data from the final trial are shown in Figure 3. These are referred to as Ugi1, Ugi3, and Ugi4, with Ugi standing for *Utricularia gibba* insulator. Ugi1 showed GFP levels comparable to the EXOB insulator (*p*= 1.5E-08). The insulator activity of Ugi4 is the strongest one we have observed so far (*p*= 2.7E-08).

**Fig 3.**
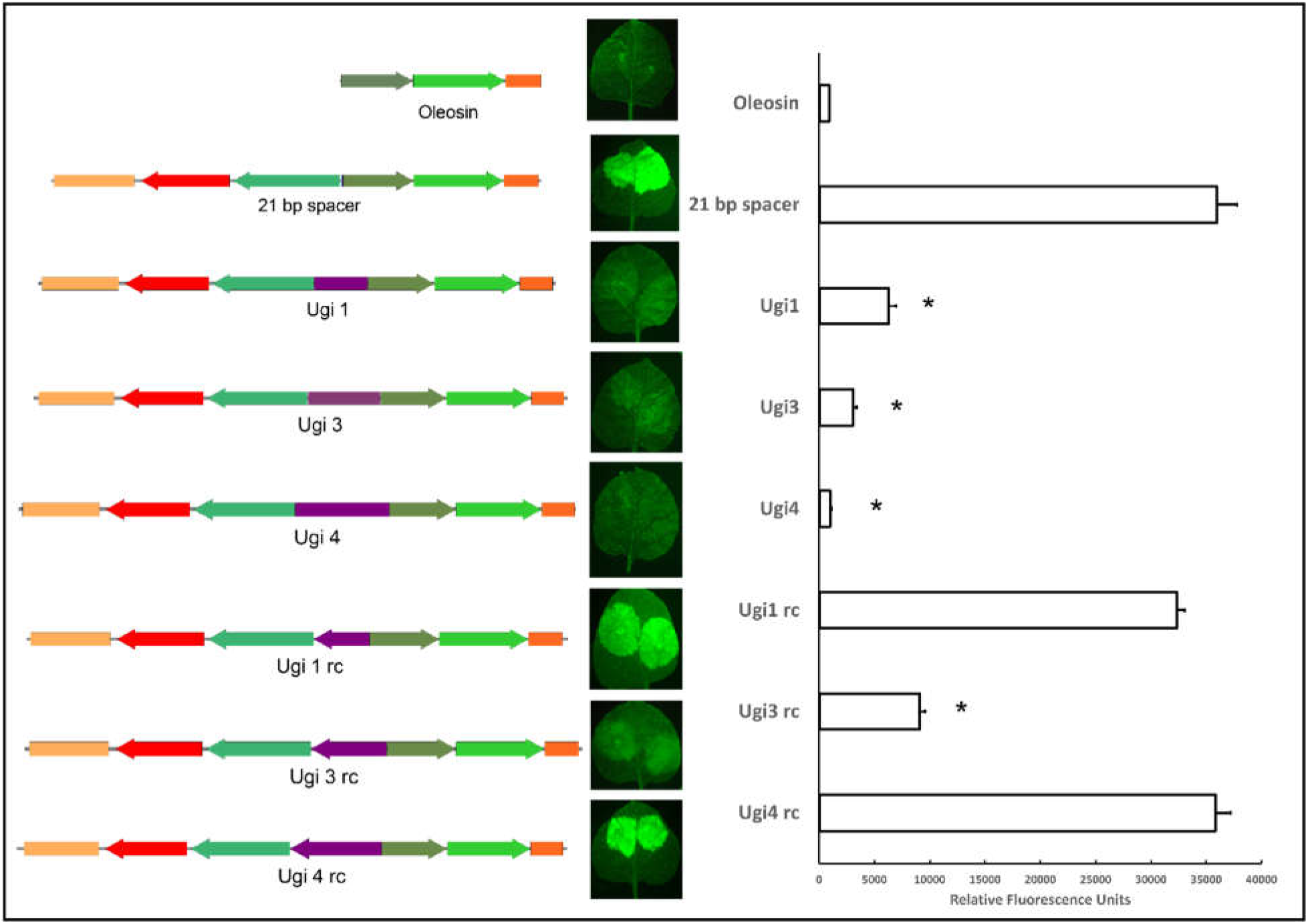
Bladderwort insulators in forward and reverse (rc) orientation. Components are as described for Figure 1. *Indicates t-test significant difference from the control construct (Oleosin promoter with 21 bp between cassettes) p < 0.001. Error bars represent standard error of the mean.

The function of some animal insulators and of the TBS insulator is orientation-dependent and therefore is more effective in one orientation than when reversed [26, 27]. To test if this is also the case with the identified bladderwort insulators, the reverse complements of Ugi1, Ugi3, and Ugi4 were evaluated. Insulator activity as shown by GFP expression was reduced for all three insulators, though the reverse complement of Ugi3 still reduced ectopic GFP expression relative to that of the control (Figure 3).

Bladderwort insulators were compared with currently available insulators for plants (Figure 4). The TBS insulator showed high levels of ectopic GFP expression, meaning it has poor insulator activity in our assays. In contrast, EXOB is four times more effective than TBS, despite being half the size, which makes it comparable to Ugi 1. The HIV-LTR sequence also showed insulator activity, though not as effectively as that of EXOB or the Ugi insulators.

BEAD1c showed a significantly lower level of GFP expression compared that of the 21-bp spacer control, but still more than double that of the EXOB insulator. GFP expression from the UASrpg construct showed no significant difference compared to that of the 21-bp control, so it has no insulator activity in our test system (Fig 5).

BEAD1c and UASrpg are described as being context-dependent [28]. For insulators to be useful in biotechnology, they need to function in a variety of genic contexts. Hence, we tested our bladderwort insulators with a second tissue-specific promoter: the APETALA3 (AP3) flower-specific promoter from *Arabidopsis* [29]. The AP3 500-bp spacer construct showed higher ectopic GFP expression than the AP3 21-bp spacer construct. Ugi1 did not show significant differences in GFP expression compared to the 21-bp control (*p*= .008). For Ugi3 and Ugi4, results from this assay very closely matched the results with the oleosin promoter, with respective *p* values 1.13E^-06^ and 1.07E^-06^ (Fig. 6).

**Fig 4.**
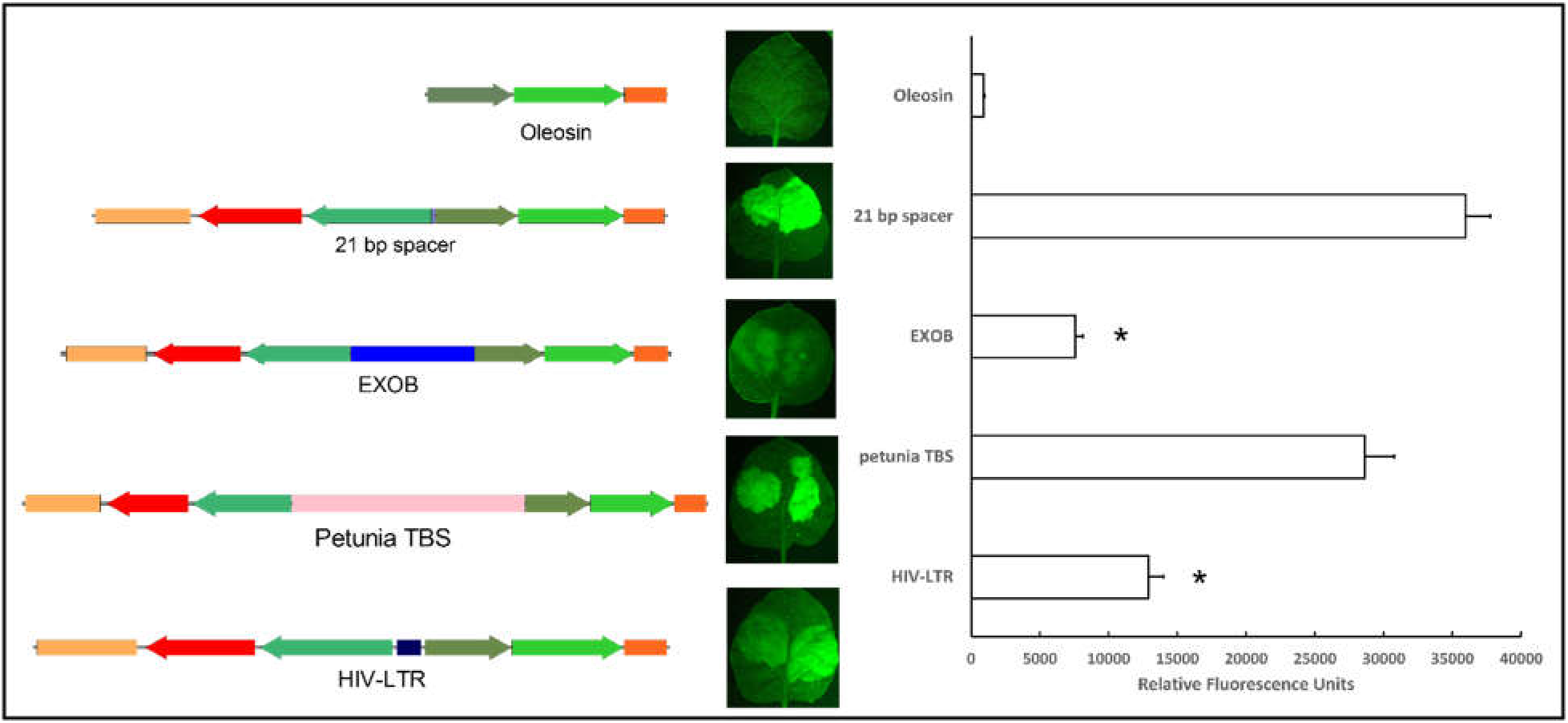
Insulators previously reported to work in plants. Components are as described for Figure 1. *Indicates t-test significant difference from the control construct (Oleosin promoter with 21 bp between cassettes) p < 0.001. Error bars represent standard error of the mean

**Fig 5.**
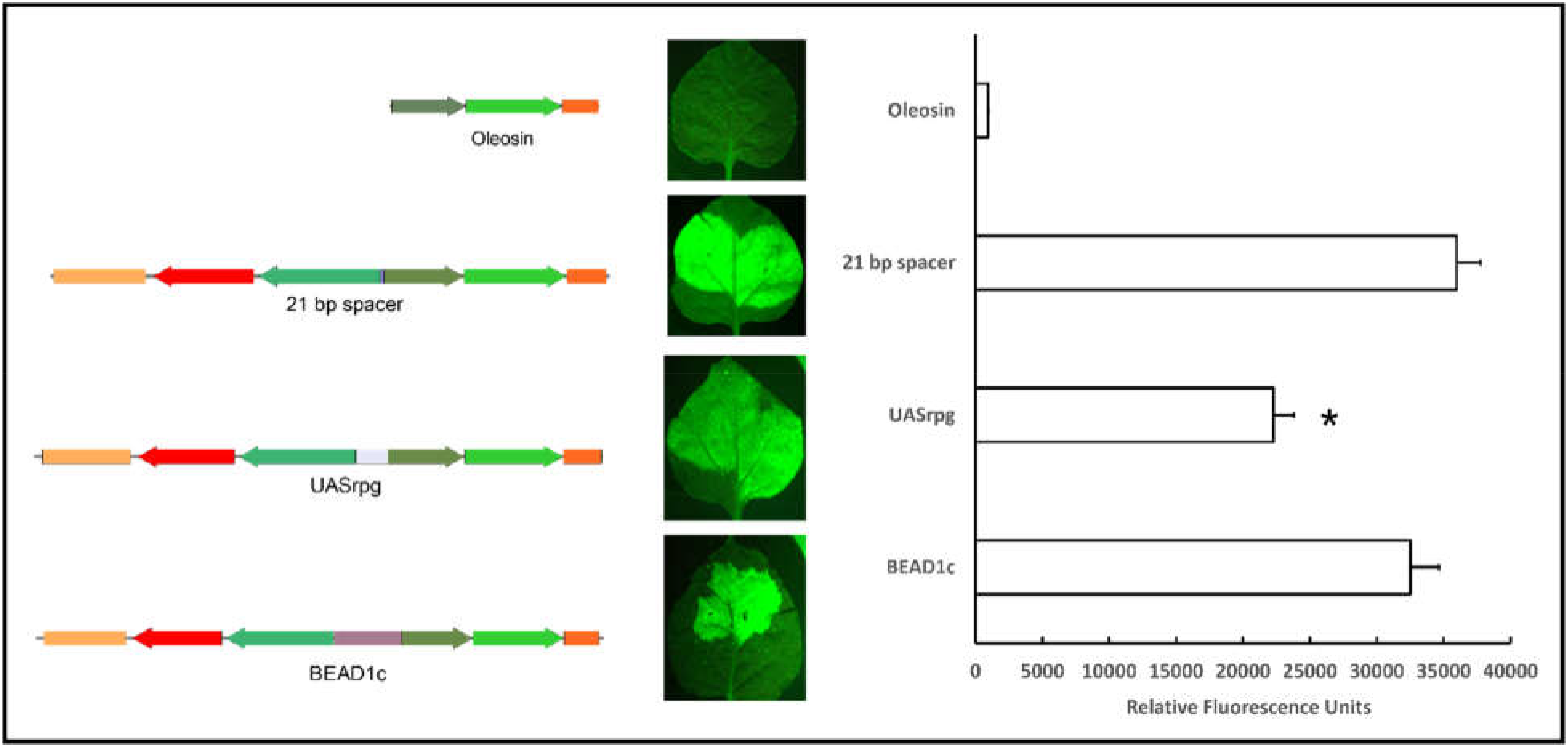
Fungal and human insulators UASrpg and BEAD1c. Components are as described for Figure 1. *Indicates t-test significant difference from the control construct (Oleosin promoter with 21 bp between cassettes) p < 0.001. Error bars represent standard error of the mean.

**Fig 6.**
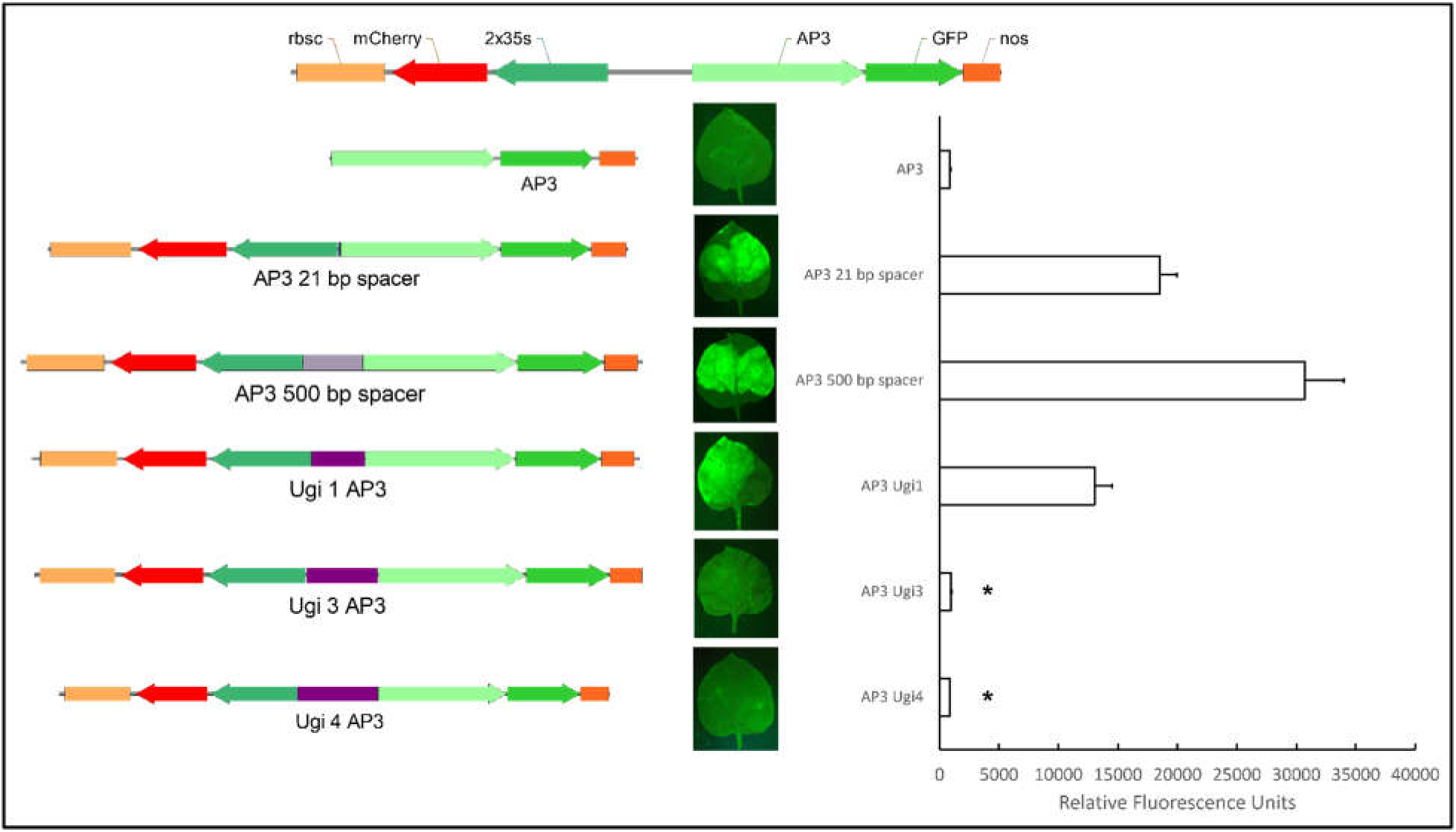
Bladderwort insulators with the flower-specific Apetala 3 promoter. rbsc: rubisco small subunit terminator from *Pisum sativum*, 2×35S: duplicated 35S promoter, AP3: *Arabidopsis thaliana* Apetala 3 promoter, nos: Nopaline synthase terminator. *Indicates t-test significant difference from the control construct (AP3 promoter with 21 bp between cassettes) p < 0.001. Error bars represent standard error of the mean.

The silencing ability of insulators in plants has not been sufficiently evaluated. All of the sequences evaluated in this study that showed insulator activity, including Ugi1, Ugi2, and Ugi4, significantly decreased not only GFP expression, but mCherry expression as well, with respective p values 1.92E-07, 4.32E-07, and 2.40E-07 (Fig. 7), pointing to the idea that these sequences may be partially silencing the flanking 35S promoter and may better be classified as silencers rather than insulators. While other studies have used a similar construct design to evaluate insulator activity [14], few have described or adequately tested for the effect that the intervening sequence has on the gene driven by 35S. Savitch et al. (2009) tested various sequences with the 35S promoter driving hygromycin phosphotransferase, which they refer to as hptII [14]. Transformed plants were selected on medium supplemented with hygromycin, but hygromycin resistance or hptII protein expression was not quantified. Other studies utilized a 35S:GFP cassette similar to our 2×35S:mCherry cassette, but did not evaluate GFP expression in any form [12, 30]. Yet other studies using a 35S:GFP cassette confirmed GFP expression via fluorescent imaging, but did not quantify fluorescence or expression [31]. Only one study evaluating the effect of a gypsy-like sequence from Arabidopsis as an insulator evaluated the gene driven by 35S through real-time RT-PCR and found no significant differences in expression, indicating that silencing was not occurring [13]. True enhancer-blocking insulators can block enhancer-promoter interference without affecting the expression flanking genes, so the evaluation of the gene driven by the enhancer-containing promoter is critical in identifying true insulators.

**Figure 7.**
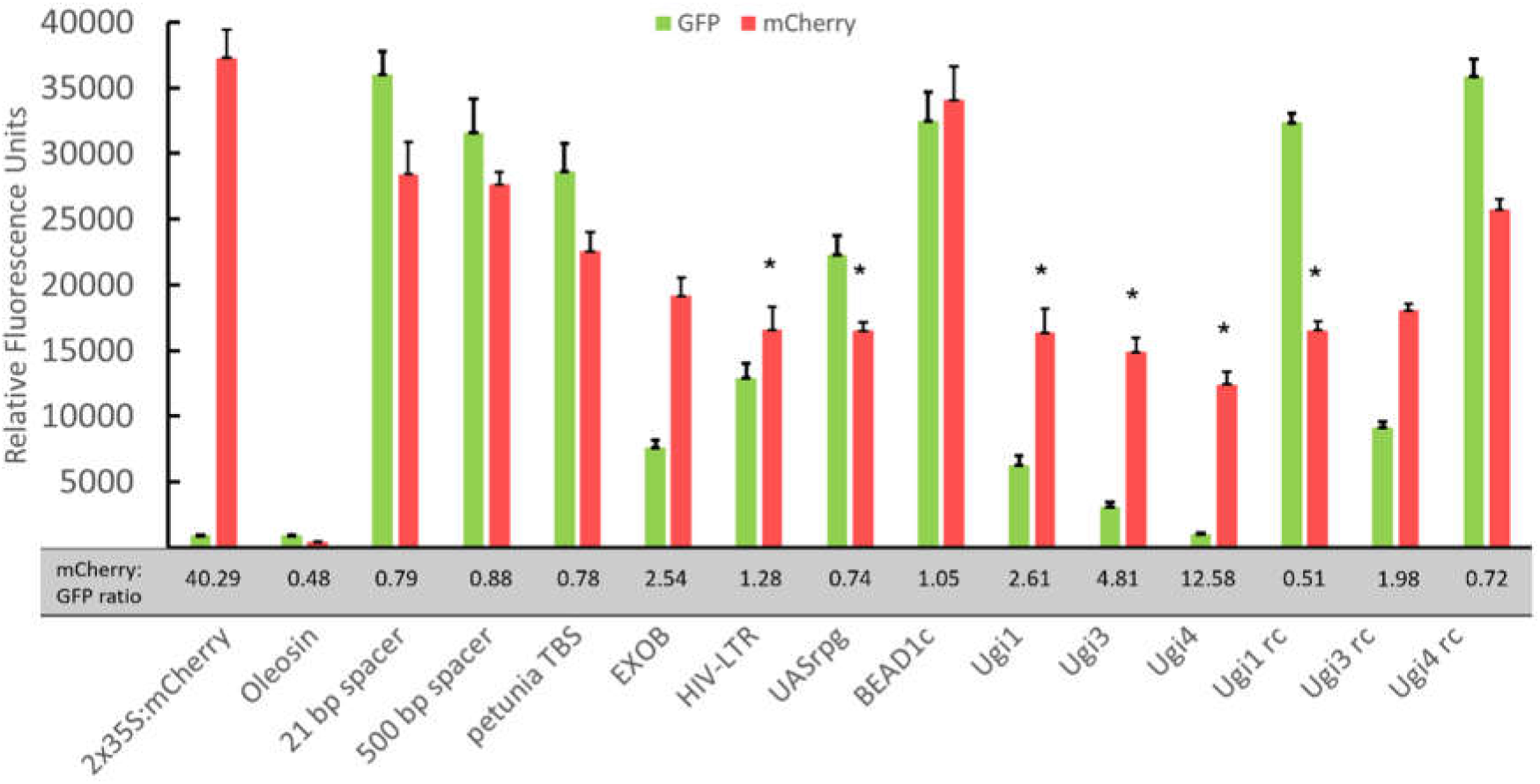
Effect of putative insulator sequences on the expression of 2×35S:mCherry. *Indicates t-test significant difference from the control construct (Oleosin promoter with 21 bp between cassettes) p < 0.001. Error bars represent standard error of the mean.

## Discussion

Variable transgene expression has been an issue with plant transformation since the beginning, though the reason has not always been clear. Since the effect of genomic insertion site on transgene expression is minor among independent, single-copy transgenics [32], any differences in expression probably come from endogenous enhancers in the genome, rather than from complex T-DNA arrangements at the insertion site [6, 32].

In some cases, interference can be attenuated by spacer DNA. In our assays, 500 bp of spacer DNA did not significantly reduce ectopic GFP expression compared to only 21 bp of spacer DNA. In contrast, kilobases of spacer DNA [14] have reduced interference; however, the repeated use of such long pieces of spacer DNA is unrealistic in multigene constructs as transformation efficiency at both the bacterial and plant level is hindered by large cassette sizes [33]. This limitation on vector size dictates that genes be closely spaced to each other, increasing the chance of intergenic interference [34].

The alternative is to use insulators to overcome interference with gene expression, so long as the insulators do not overly inflate the size of the resulting vectors. To find appropriate insulators, this work used a system of vectors very similar to those used previously to identify insulator function [11, 14], but our work differs in several key aspects. First, we devised a bioinformatics screen based on gene expression, closeness of gene pairs, and conservation across asterids to identify sequences with putative insulator function. Secondly, agroinfiltration (the method used in the current study) provides feedback from millions of individually transformed cells, while stably transformed plants are derived from single transformed cells that, depending on the transgene insertion site, can exhibit transcriptional interference from endogenous enhancers. However, our results reflect expression from both integrated and non-integrated T-DNA at three days after infiltration. It is not known if insulators differently affect genes that are not yet integrated into the genome.

Of the 43 putative insulators found, the most promising 10 were chosen for *in planta* validation based on the filtering criteria: degree of expression difference between genes and the length of the putative insulator sequence. Within these candidates, four out of the 10 contain conserved sequences (30-60 bp) based on whole genome alignment with asterids. However, of these, only Ugi4 has significant insulator activity.

Looking at the top 10 identified putative bladderwort insulators from the list of 43, two (Ugi3, Ugi4) show significant insulator activity, reducing ectopic GFP expression significantly compared to the control 21-bp spacer construct, with Ugi4 nearly eliminating GFP expression. A third potential insulator, Ugi1, is effective with the oleosin promoter, but not the apetala promoter. The Ugi insulators have the added benefit of being shorter: 451, 619, and 790-bp long, respectively, for Ugi1, Ugi3, and Ugi4. All are smaller than the EXOB insulator, though still longer than desirable. Further analysis can determine which part of the sequence is needed for effective insulator activity in the Ugi insulators.

Collectively, these results indicate that these newly identified *U. gibba* sequences can function as insulators in the sense that they prevent the enhancer in the 2×35S promoter from leading to ectopic expression of GFP in the adjoining cassette. Based on the results in this test system, these Ugi insulators, along with EXOB, may currently be the most effective enhancer-blocking elements for use with plants. However, all these sequences also have silencing activity such that the expression of the 2×35S:mCherry construct is attenuated. They do stand apart in that they have more enhancer-blocking insulator activity than silencing activity than all other tested sequences except EXOB (Figure 7). Hence, they could still have utility in transgenic applications.

Prior to this work, only three plant-derived enhancer-blocking insulators had been reported. Of these, the N129 from *Arabidopsis* is reported to be undependable [14], and the petunia TBS and gypsy-like sequence from *Arabidopsis* are over 2-kb and therefore are too large to be practical in multigene-containing vectors. The petunia TBS is also among the least effective of the insulators tested here. EXOB is the more effective of the pre-existing insulators, while HIV-LTR is intermediate in effectiveness to TBS and EXOB. UASrpg is intermediate to TBS and HIV-LTR in terms of insulator activity.

Ultimately, it is still not clear what constitutes an insulator sequence. CTCF binding sites are components of insulators in animals and are numerous in the EXOB insulator, pointing to their conserved function in plants. However, CTCFs are unique to bilateran phyla [35] and plant sequences without insulation ability also have CTCF biding sites [11], so even though these sequences exist, they are likely not functional in the same way they are in animals and fungi. Likewise, the gypsy insulator from *Drosophila melanogaster* contains several Su(Hw) protein binding sites. *Arabidopsis* lacks the Su(Hw) proteins needed to bind to the gypsy insulator, but insulation was still observed by She et al. (2010) [36], even when Su(Hw) proteins were not co-expressed in the plant, suggesting that an *Arabidopsis* protein was binding to the gypsy insulator and providing insulator activity; however, another study was unable to replicate these results [16].

The observation that insulators previously reported to be effective in plants were not effective in our test system is consistent with the observation that some insulators are context-dependent and are more effective with some genes than others. Furthermore, the same element may function as an insulator, enhancer, or silencer depending on which transcription factors are expressed in any given tissue [37-39]. Therefore, it may be necessary to identify a library of gene- and tissue-specific insulator-like sequences for use in plant transformation.

Given the apparent lack of canonical insulator sequences in plants, plant sequences are better referred to as ‘insulator-like.’ Regardless, the Ugi sequences identified here can be used as insulators for plant transformation, despite their silencing activity, if moderate expression levels of the flanking gene are sufficient to meet the transformation objectives.

## Methods

### Plant material

*U. gibba* plugs were obtained from Meadowview Biological Research Station (https://www.pitcherplant.org) and were propagated in three separate tanks. Plugs were anchored in a layer of sand above a layer of peat moss under approximately six inches of municipal water. Illumination was provided by 50-watt LED grow lights (Morsen) were placed outside each tank on a timer of 16 hours on / 8 hours off. Water was changed and the tanks cleaned as needed.

### Whole-genome sequencing

DNA from bulk bladderwort tissue was isolated using the Promega Wizard Genomic DNA Purification Kit. Libraries were prepped using a KAPA library prep kit (#KK8231) and sequenced on two Illumina MiSeq flowcells with paired-end 300-bp reads at the Georgia Genomics and Bioinformatics Core.

### RNA sequencing

Live bladderwort was placed in ice water and dissected under the microscope to obtain 10 cm each of rhizoids and stems, 30 leaves, and 30 bladders from each of three biological replicates. RNA was isolated from these individual tissues and from whole plants using TRIzol/chloroform extraction (Invitrogen). Full-length RNA transcripts were prepared from bulk tissue using a stranded KAPA kit (#KK8420) and sequenced on two Illumina MiSeq flowcells, producing paired-end 250-bp reads. Tissue-specific libraries of 3’ RNA transcripts were prepared using a Lexogen QuantSeq 3’ mRNASeq FWD kit and sequenced on a mid-output NextSeq 500 flowcell, producing single-end 150-bp reads.

### Genome assembly and annotation

Raw paired-end genomic reads were merged and trimmed in Trimmomatic v0.36 using the “ILLUMINACLIP” option with default parameters [40], keeping only trimmed reads 36 bp or longer. Read quality was evaluated with FastQC v1.8.0 [41] and MultiQC v1.5 [42]. Genome assembly was performed with SPAdes v3.13.1 [43] with default parameters, and quality metrics determined with QUAST v5.0.2 [44]. Contigs originating from contaminant, mitochondrial, and chloroplast genomes were filtered out using Kraken v2.0.7 [45] with a database of bacterial, fungal, and plant sequences from NCBI plus published *U. gibba* genome sequences [46].

Annotation was performed in multiple rounds using Maker v2.31.10 [47] First, basic annotation was performed using Trinity v2.6.6 [48] based on whole-transcript RNA-Seq and protein models from four other sequenced asterid genomes: *Daucus carota (GCF_001625215*.*1), Helianthus annuus (*GCF_002127325.1*), Nicotiana attenuate (*GCF_001879085.1), and *Solanum lycopersicum (*GCF_000188115.4). After the initial round of annotation, putative genes were pulled out and used to train a gene prediction model in Augustus v3.2.3 [49]. Gene models were refined in the next rounds of annotation using Augustus gene prediction models trained after each Maker run. Maker was run for three rounds, at which point the Annotation Edit Distance (AED) score distribution stopped improving. Finally, the published PacBio *U. gibba* assembly [20] was used to BLAST against the annotated genome to identify any proteins the pipeline had missed. Only BLAST results with at least 95% sequence similarity and 90% sequence length that did not overlap an existing Maker annotation were kept. Overlaps were found using BedTools intersect v2.29.2, [50].

### Mining for regulatory regions using 3’ RNA-Seq data

Raw 3’ RNA-Seq reads were trimmed using Trimmomatic v0.36 using the “ILLUMINACLIP” option, with max adapter mismatch count equal to two, palindrome clip threshold equal to 30, and match accuracy to 10. Quality metrics were then visualized using FastQC v1.8.0 and MultiQC v1.5 [41, 42]. Reads were mapped to the genome using the STAR aligner v2.7.0 [51] and reads per gene were quantified using htseq-count v0.9.1 [52]. Once counts were obtained, they were normalized in DESeq2 using the standard “DESeq” function [53].

Pairs of genes were extracted and classified as either “divergent,” “convergent,” or “parallel” based on their relative orientations. Expression levels quantified as the relative expression difference between the two genes (“relative fold difference”). Intergenic regions were selected as potential insulators if the flanking genes (1) were both expressed, (2) were in the top 10% of gene pairs based on their relative fold difference, (3) were <1000 bp apart, and (4) the higher-expressed gene was in the top 50% of expressed genes in the dataset. Once candidate regions meeting these criteria were selected, they were prioritized by relative fold change, sequence length (shorter preferred over longer), and lack of correlation of neighboring gene expression across the tissue-specific datasets.

### Intergenic sequence conservation and presence of putative insulator sequences

Conserved intergenic sequences were found using whole genome multiple alignments of different clades of angiosperms to the newly assembled *U. gibba* genome. The four clades comprising whole genome alignments were asterids (*Vaccinium corymbosum, Actinidia chinensis, Lactuca sativa, Helianthus annuus, Cuscuta australis, Capsicum annuum, Solanum lycopersicum, Petunia axillaris, Coffea canephora, Utricularia gibba, Mimulus guttatus, Genlisea aurea*), rosids (*Kalanchoe fedtschenkoi, Carica papaya, Arabidopsis thaliana, Durio zibethinus, Cannabis sativa, Prunus persica, Glycine max, Medicago truncatula, Betula pendula, Cirtrullus lanatus, Manihot esculenta, Vitis vinifera*), and monocots (*Zostera marina, Spirodela polyrhiza, Ananas comosus, Setaria italica, Sorghum bicolor, Brachypodium distachyon, Oryza sativa, Asparagus officinalis, Phalaenopsis equestris*). Alignments of each clade to *Utricularia gibba* were completed according to the UCSC genome browser full genome alignment tutorial (http://genomewiki.ucsc.edu/index.php/Whole_genome_alignment_howto), but using LastZ v1.04.03 [54] instead of MultiZ for the initial alignment step. After alignments were completed and combined, PhastCons v1.5 [55] was used to generate genome-wide sets of conserved coordinates for each clade with options target-coverage = 0.125 and expectedlength = 20; all other options were set to default. Analysis of conserved region distribution in gene pairs was carried out with custom R scripts.

### Entry vector assembly and transformation

Entry vectors for Golden Gate cloning were made by digesting the Greengate Kit [56] entry vector plasmids (PGGA000, PGGB000, PGGC000, PGGD000, PGGE000, PGGF000) with BSA1-HF V2 (New England Biolabs Inc.). Bladderwort sequences were synthesized (IDT, Twist, Genewiz) and amplified by PCR (Table S3) to create 20-bp of homology with the digested vector on each side. The reaction leaves 21 bp between cassettes. In addition, a 500-bp spacer DNA fragment (Table S4) was randomly generated with the Random DNA Sequence Generator (http://www.faculty.ucr.edu/~mmaduro/random.htm).

Gibson assembly was performed by combining 1 μL of digested vector, 1 μL of gel-purified PCR product, and 2 μL of NEBuilder Hifi DNA Assembly Mastermix 2x and incubating at 50°C for 15 minutes. *E. coli* transformation was performed with DH5α chemically competent cells following the protocol from New England Biolabs. Transformants were screened for the insert sequence with PCR. Positive transformant colonies were cultured overnight and purified using the Epoch GenCatch Plasmid DNA Mini-prep kit. Completed entry vector plasmids were confirmed by Sanger sequencing (Genewiz).

### Golden Gate assembly and transformation

Completed entry vector plasmid stocks were quantified by A260 absorbance and diluted to 100 ng μL^-1^. Golden Gate cloning was performed as described by Decaestecker et al. [57]. The assembled constructs and their components are listed in Table S2. One μL of finished reaction product was transformed into 20 μL *E. coli* competent cells (genotype DH5α) following the supplied protocol and plated on Luria Bertani (LB) agar plates supplemented with spectinomycin (100 μg mL^-1^) to select for positive transformants. Single positive transformant colonies were cultured overnight and purified using the Epoch GenCatch Plasmid DNA Mini-prep kit. Completed entry vector plasmids were confirmed by Sanger sequencing. The destination vector used for every Golden Gate assembly reaction was pGGP-AG-nptII, created by inserting the Stubi3P:nptII:Stubi3T cassette from AddGene plasmid 59175 between the XbaI/KpnI sites of pGGP-A-G [57].

### *Agrobacterium* electroporation

Twenty-five μL of electrocompetent *Agrobacterium tumefaciens* strain LBA4404 cells were electroporated in an electroporator (Bio-Rad Micropulser) according to the manufacturer’s instructions with 1 μL of Golden Gate assembled plasmid prep.

### *Nicotiana* agroinfiltration

*Nicotiana benthamiana* TW17 plants were grown in 2.54-cm (2 inch) pots using SunGro potting mix. Plants were grown under a 12:12 hour light/dark photoperiod at 22°C and 24°C respectively. Agroinfiltration of 4-week-old plants was carried out as described by Felippes and Weigel 2010 [58]. Twenty mL of LB liquid medium with the appropriate antibiotics were inoculated with a single colony of *A. tumefaciens* and incubated for 48 hours shaking at 28°C. After incubation, cells were pelleted, supernatant was removed, and the cells were resuspended in infiltration buffer (10 mM MgCl2, 10 mM MES, 100 μM acetosyringone) and diluted to an optical density of 0.75 and incubated at room temperature for 3 hours. Three leaves in each of three plants were infiltrated per construct with a 1-mL needleless syringe. After infiltration, plants were kept for 3 days at 12-hr daylight, 24°C day, 22°C night temperatures. The Xite Fluorescent Flashlight system (Nightsea, Designation RB-GO) with excitation at 440-460 nm and emission at 500-560 nm was used to visualize GFP in leaves. Pictures were taken with a Canon EOS 60D camera fitted with a green camera lens (Tiffen Green #58), manual settings, a 6-second exposure time, and aperture 2.8.

### Fluorescence microtiter plate spectroscopy

Fluorescence microtiter plate spectroscopy of intact *Nicotiana benthamiana* leaf discs was performed as described by [59]. Single leaf discs were obtained with a 6-mm diameter cork borer and floated adaxial side down in the wells of an opaque 96-well plate with 300 μL of water. Fluorescence detection was performed using a Synergy 2 reader with Gen5 version 3.09 software (BioTek Instruments Inc., Winooski, VT, USA) with excitation at 475 - 495 nm, and emission at 508-548 nm for GFP and excitation of 587 nm and emission of 610 nm for mCherry. For each construct, a total of nine leaf discs were sampled across three infiltrated plants. Three leaves were infiltrated per plant, and one leaf disc was taken from each.

### Experimental design and statistical analysis

For each treatment there were three biological replicates (three plants) and three technical replicates (three leaf discs). Three leaves were infiltrated per plant, and one leaf disc was sampled from each leaf, resulting in nine leaf discs per treatment. Raw fluorescence reads were compared by analysis of variance (ANOVA) followed by two-sample t-tests between the Oleosin 21 bp spacer or AP3 21 bp spacer controls and the bladderwort insulator outputs to determine significant differences in GFP expression. All statistically significant results were judged based on a *p* value of ≤.001.

### Materials availability

Ugi1, Ugi2, and Ugi3 are available as GenBank OK086967, OK086968, and OK086969, respectively. Plasmids with the Ugi sequences are also publicly available on AddGene (ID numbers 177194, 177195, 177196)]

All bioinformatic scripts for this analysis, including assembly and annotation, are available at https://github.com/lkov0/bladderwort-analysis. Raw DNA, RNA, and 3’ RNA reads are available at the NCBI Sequence Read Archive (BioProject PRJNA595351).

## Acknowledgements

This work was funded by the National Science Foundation, grant IOS-1759827. We thank Holly Griffis, Orrie Fetisma, Erik Schouten, and Zachary Carter for their excellent technical assistance. Finally, we thank Bob Schmitz for helpful discussions on the interpretation of the results.

## Author contributions

LK: Carried out bioinformatic analyses; wrote & edited manuscript

PRL: Designed and help build all the vectors for the project

JP: Performed molecular cloning and *in planta* assays; wrote & edited manuscript

WAP: Conceived the project & initial experimental design, supervised execution; edited manuscript

JGW: Designed experiments; supervised analyses; edited manuscript

## Supplementary material

**Table S1:** Bladderwort insulator candidates.

| Bladderwort insulator | Length (bp) |
| --- | --- |
| Ugi1 | 451 |
| Ugi2 | 23 |
| Ugi3 | 619 |
| Ugi4 | 790 |
| Ugi6 | 124 |
| Ugi7 | 194 |
| Ugi11 | 342 |
| Ugi14 | 74 |
| Ugi15 | 258 |
| Ugi20 | 24 |

**Table S2:** Genome assembly metrics.

|  | Reference Genome [20] | PP01 draft assembly |
| --- | --- | --- |
| Length | 102Mb | 101Mb |
| # Contigs | 581 | 13033 |
| # Genes | 29,666 | 27,166 |
| BUSCO % | 88% | 84% |

**Table S3:**
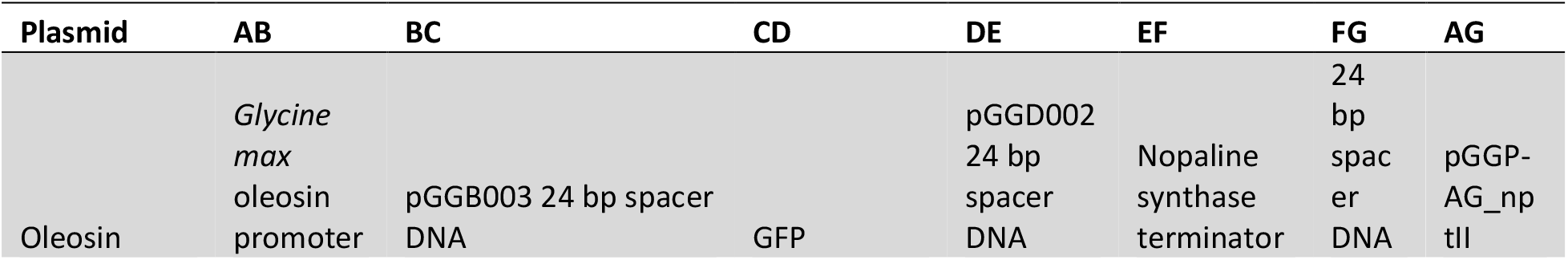
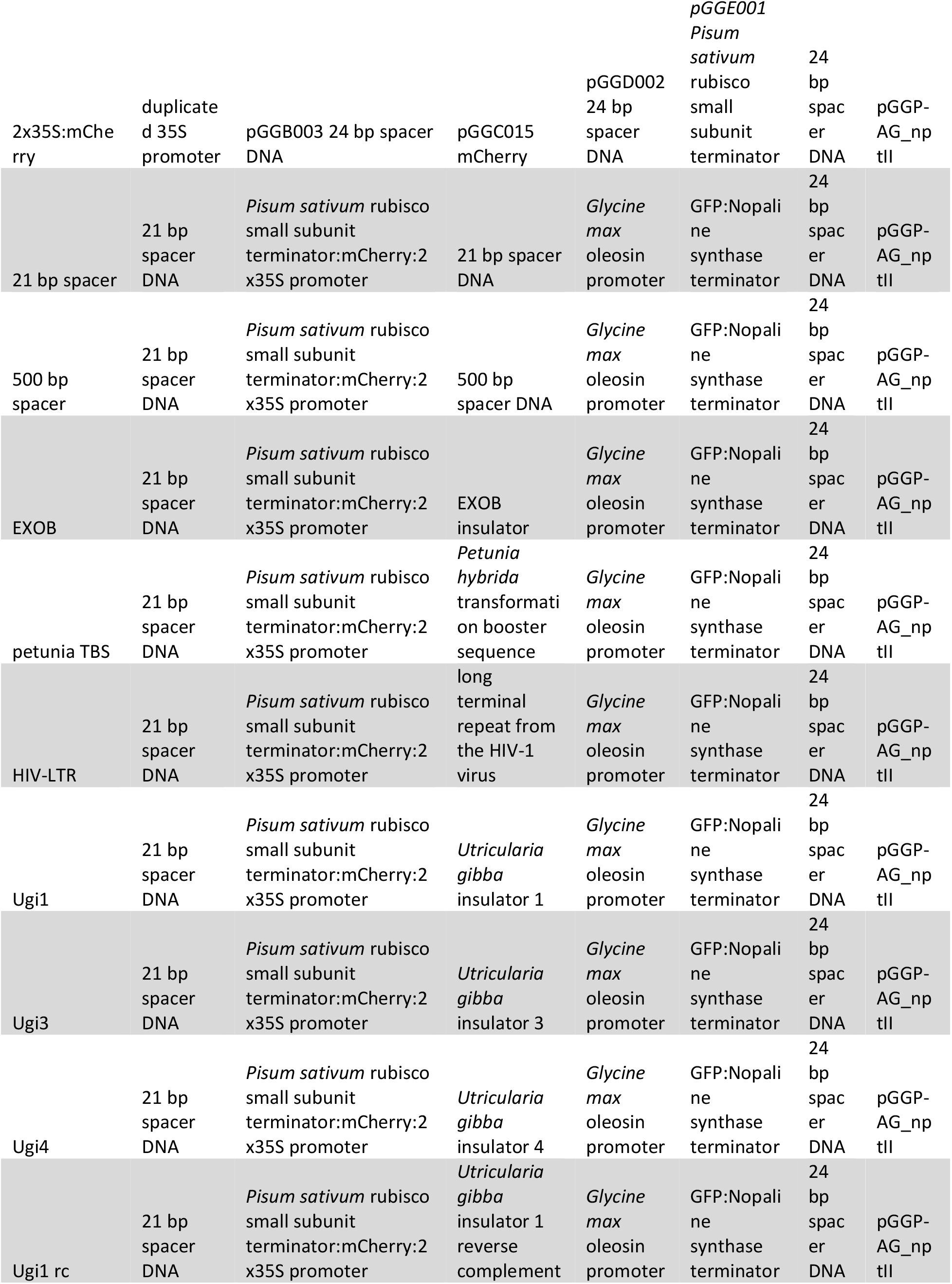
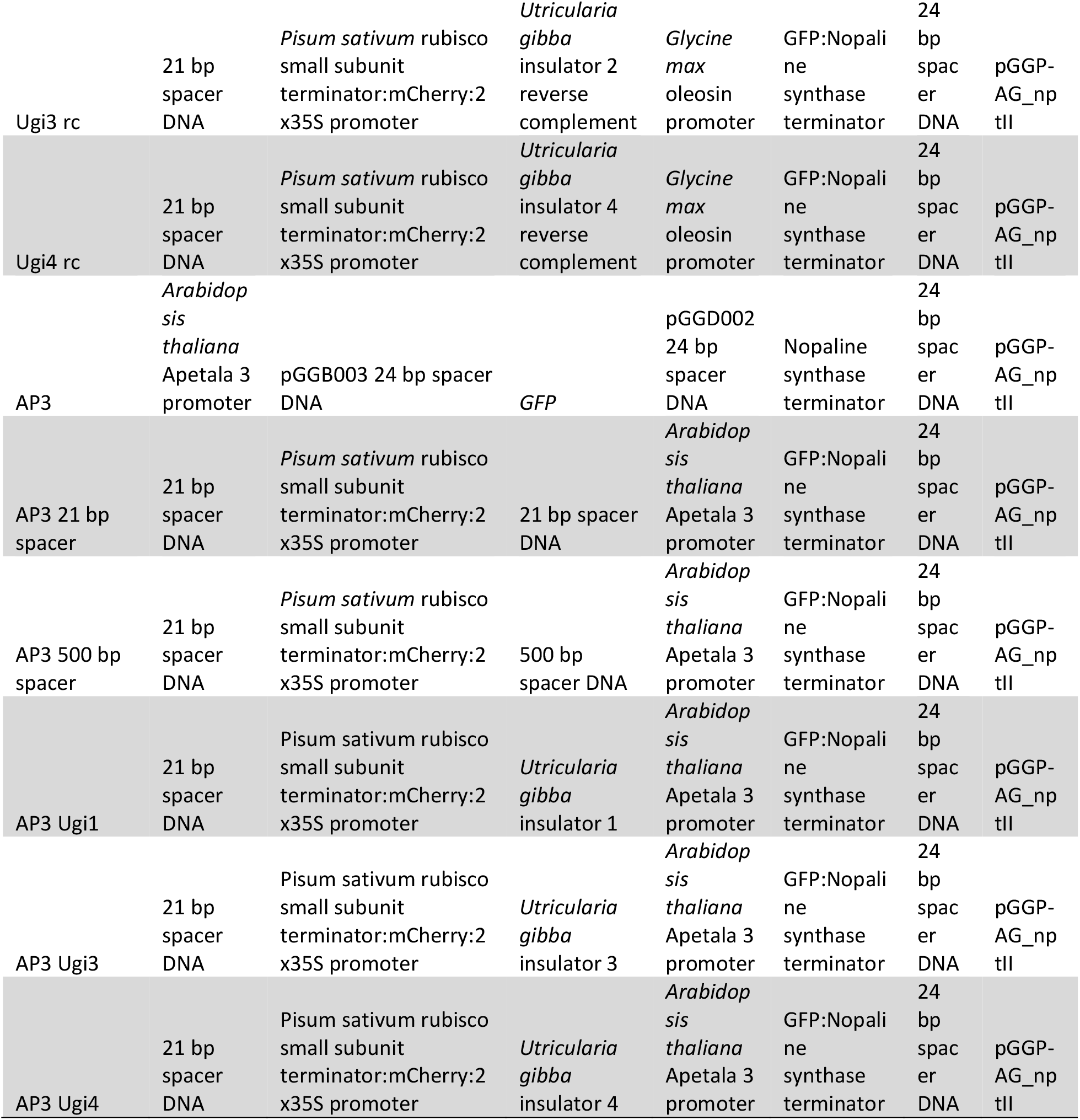
Vector components.

**Table S3:**
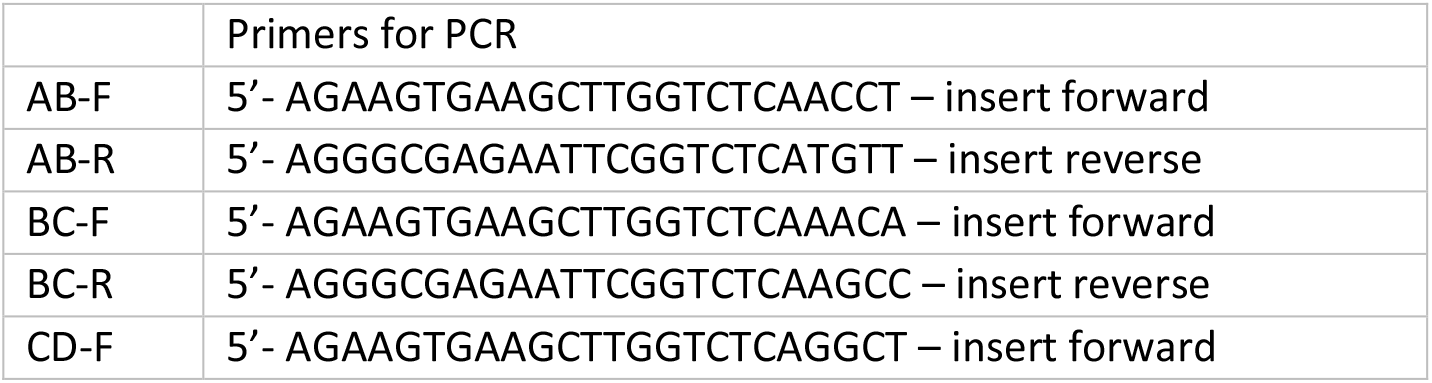
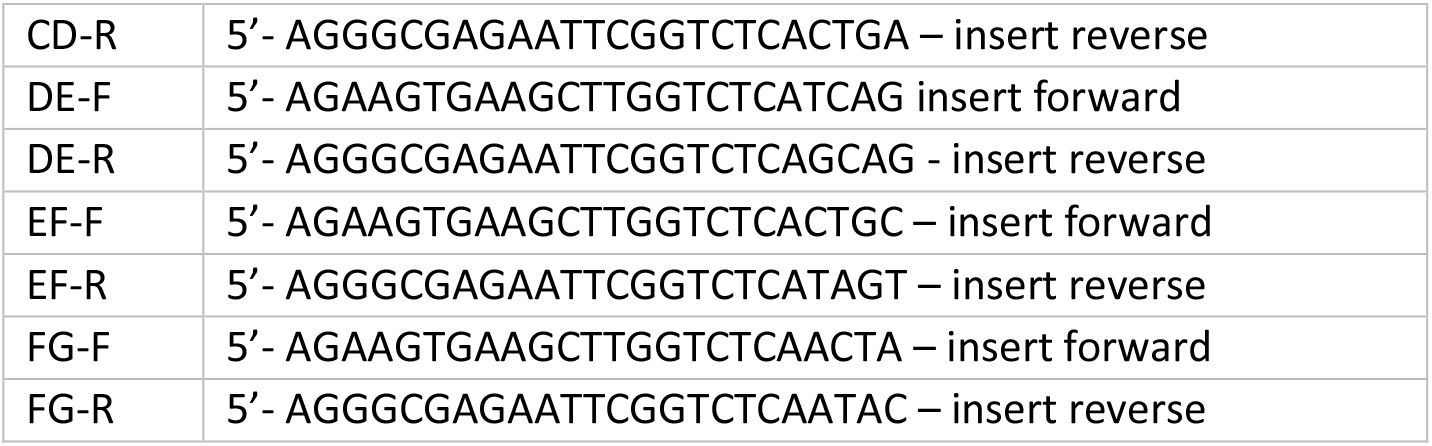
Entry vector primer sequences.

**Table S4:**
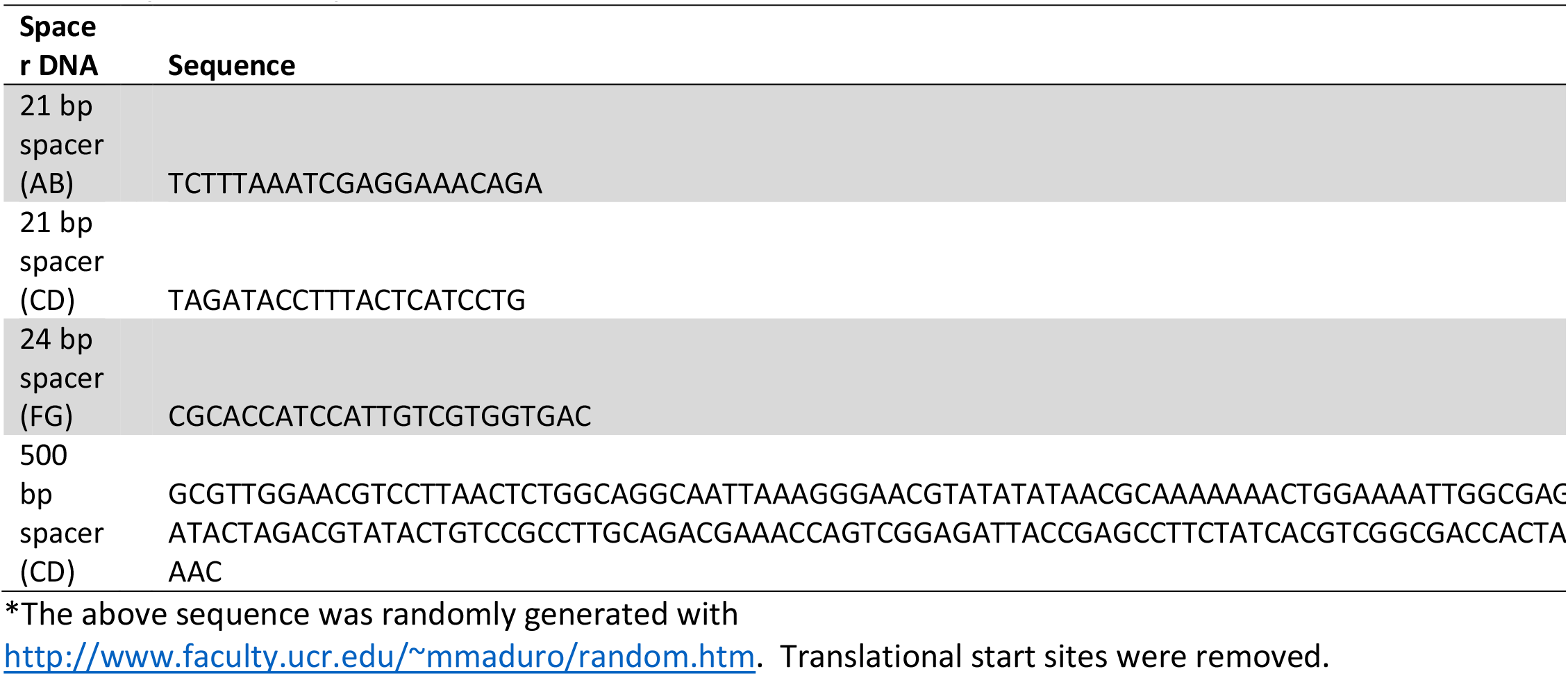
Spacer DNA sequences *.

